# Comparing 10x Genomics single-cell 3’ and 5’ assay in short-and long-read sequencing

**DOI:** 10.1101/2022.10.27.514084

**Authors:** Justine Hsu, Julien Jarroux, Anoushka Joglekar, Juan P. Romero, Corey Nemec, Daniel Reyes, Ariel Royall, Yi He, Natan Belchikov, Kirby Leo, Sarah E.B. Taylor, Hagen U Tilgner

**Affiliations:** Feil Family Brain and Mind Research Institute, Weill Cornell Medicine, New York, NY, USA; Center for Neurogenetics, Weill Cornell Medicine, New York, NY, USA; 10x Genomics, Pleasanton, CA, USA; Physiology, Biophysics & Systems Biology Program, Weill Cornell Medicine, New York, NY, USA

**Author notes:** corresponding author: Hagen U Tilgner. equal contrition (Justine Hsu, Julien Jarroux).

**Keywords:** Single-cell, Long-read, Short-read, Technology comparison

## Abstract

Barcoding strategies are fundamental to droplet-based single-cell sequencing, and understanding the biases and caveats between approaches is essential. Here, we comprehensively evaluated both short and long reads of the cDNA obtained through the two marketed approaches from 10x Genomics, the “3’ assay” and the “5’ assay”, which attach barcodes at different ends of the mRNA molecule. Although the barcode detection, cell-type identification, and gene expression profile are similar in both assays, the 5’ assay captured more exonic molecules and fewer intronic molecules compared to the 3’ assay. We found that 13.7% of genes sequenced have longer average read lengths and are more complete (spanning both polyA-site and TSS) in the long reads from the 5’ assay compared to the 3’ assay. These genes are characterized by long average transcript length, high intron number, and low expression overall. Despite these differences, cell-type-specific isoform profiles observed from the two assays remain highly correlated. This study provides a benchmark for choosing the single-cell assay for the intended research question, and insights regarding platform-specific biases to be mindful of when analyzing data, particularly across samples and technologies.

## Introduction

The development of single-cell genomics (Klein et al., 2015; Macosko et al., 2015; Zeisel et al., 2015) has allowed a deeper analysis of gene expression across the cell types that form complex tissues such as the brain (Network, 2021; Yao et al., 2021), blood (Villani et al., 2017; Wang et al., 2021) and lung (Habermann et al., 2020; Travaglini et al., 2020). Given the advances in long-read sequencing (Au, 2022; Hardwick et al., 2019), we and others have devised long-read methods to measure isoform expression through TSS, splice sites, polyA sites, and full isoform sequencing across thousands of single cells (Gupta et al., 2018; Joglekar et al., 2021; Singh et al., 2019; Volden & Vollmers, 2022) and nuclei (Hardwick et al., 2022). Fundamental to these approaches are the barcoding strategies which often employ microfluidics, most notably 10x Genomics. This company currently has two marketed approaches related to gene expression; the “3’ assay” and the “5’ assay”. The two assays capture different ends of the transcript in the final short-read sequencing library. Both workflows use poly-dT primer for reverse transcription and a template-switching oligo (TSO) to reverse-transcribe and amplify full-length transcripts. In the 3’ assay, poly-dT is barcoded and further selected through ligations and PCR to capture RNA at the 3’-end (Fig. 1a). In the 5’ assay, poly-dT only serves as RT primer while the TSO holds the barcode and is selected in order to sequence the 5’-end of the RNA transcripts (Fig. 1b).

**Fig. 1:**
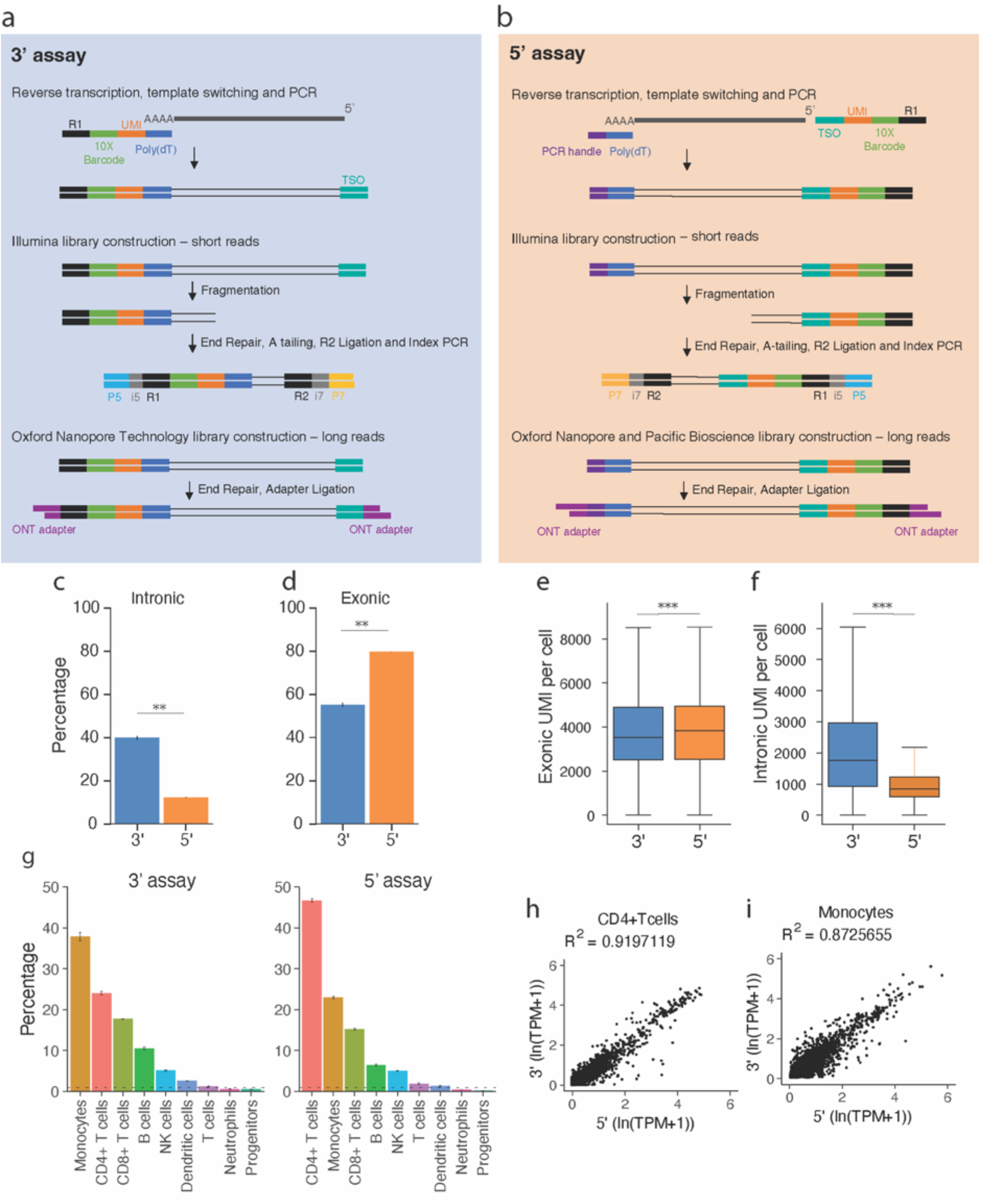
Short read analysis of single-cell 3’ and 5’ assays with PBMC data. **a**. Schematic of the molecules captured with 3’ assay for short-read and long-read sequencing. **b**. Schematic of the molecules captured with 5’ assay for short-read and long-read sequencing. **c**. Barplots showing the average percentage of mapped reads in introns and **d**. the average percentage of mapped reads in exons of two replicates from both assays. *P*-values were calculated with the Student’s t-test (***p < 10^−10^, **p < 10^−5^, *p < 0.05). **e**. Boxplots of exonic UMI per cell, **and f**. boxplots of intronic UMI per cell. Down-sampled each replicate to 10 thousand reads per cell after excluding reads with no barcode and reads for cells with less than 10 thousand reads. Combined the two replicates for each assay. *P*-values were calculated using a two-sided Wilcoxon rank-sum test (***p < 10^−10^, **p < 10^−5^, *p < 0.05). **g**. Barplots showing the percentage of cells in each cell type from each assay; Dash line denotes the 1% cut-off for cell types excluded. **h**. Scatter plot showing the gene expression level of CD4+Tcells between 3’ and 5’ assay. **i**. Scatter plot showing the gene expression level of Monocytes between 3’ and 5’ assay.

Of note, microfluidic production of cDNAs using poly(dT) priming is subject to a number of potential artifacts, including, but not limited to, priming from intronic polyA-rich regions, incomplete cDNA production, or bias against some molecules during PCR.

Given the existence of the two assays, a fundamental question is whether both reveal correlated measurements or whether one might be subject to a bias that the other one may avoid. To answer this question, we sequenced two replicates from each assay to systematically evaluate the biases and artifacts of each capture technique and its consequences on long-read sequencing. We find that the 5’ assay captures longer transcripts for a group of lowly expressed genes with longer transcript length, but the general trend of cell type-specific isoforms captured remains highly correlated.

### Short read analysis of single-cell 3’ and 5’ assays with PBMC data

Identifying cell types by clustering single-cell short-read data is the first step in defining cell-type-specific traits. While the construct structure differs between the two assays, cell-type identification methods are largely similar for both approaches. Therefore, we compared single-cell short-read sequencing of two human PBMC samples (one for each assay) with four technical replicates each. Interestingly, the 5’ assay has 28% fewer mapped reads in introns and 25% more mapped reads in exons compared to the 3’ assay (Fig. 1c, d). In order to test if the two assays have differences in UMI captured per cell, we excluded reads from cells with less than 10,000 reads and down-sampled each sample to 10,000 reads per cell. Further supporting the difference in exonic and intronic mappings, we found the 5’ assay to have higher exonic UMI counts and lower intronic ones compared to the 3’ assay (p < 10^−10^, Wilcoxon rank-sum test) (Fig. 1e, f). We next identified cell types separately for each assay combining all four technical replicates. The two assays showed similar clustering profiles with the same cell types identified (Figs. 1g, S1a, b.), and we excluded cell types representing less than 1% of total cells from further analysis (Fig. 1g). We then investigated gene expression profiles of all cell types and found that gene expression of matched cell types from both assays is highly correlated (R^2^ = 0.91 for CD4+Tcells; R^2^ =0.87 for Monocytes; Fig 1h, i, and Fig S1c). The aforementioned findings were confirmed by a separate experiment where a single PBMC sample was used for both assays (Fig. S2a-d). In summary, the 3’ and 5’ assays from 10x Genomics capture gene expression profiles of different cell types without strong differences in cell-type profiling but with lower coverage of exons in the 3’ assay.

### Barcode identification of single-cell 3’ and 5’ assay long-read sequencing

Barcode identification is the basis of the single-cell analysis, especially for the study of isoform expression using long-read sequencing (Gupta et al., 2018; Hardwick et al., 2022; Joglekar et al., 2021). In order to assess if the two barcoding strategies have an impact on barcode recognition and full-length isoform capture, we sequenced the same 5’ and 3’ barcoded cDNAs as previously mentioned using Oxford Nanopore (two replicates out of the four for each assay). First, we tested if barcodes can be detected with similar efficiency in long reads by looking for adjacent sequence elements such as the polyA tail at the 3’ end or the TSO sequence at the 5’ end, next to which the 3’ and 5’ cellular barcodes are located respectively. Surprisingly, the 5’ assay showed a 25 percentage point increase in the detection of the polyA compared to the 3’ assay (Fig 2a). On the other end, the difference in the presence of the TSO was milder but significant by 13 percentage points (Fig 2b). The differences in the identification of polyA in 3’ assay and TSO in 5’ assay broadly translated to the identification of barcodes in each assay (Fig 2c), since the relative success rate of identifying a barcode given the detection of the associated polyA or TSO element was similar between assays (Fig 2d). In summary, the 5’ assay library showed higher percentages of reads with identifiable polyA, TSO, and barcode. We then further investigate the distribution of the start position of polyA / TSO and barcode in the first/last 200 bases of barcoded reads. The theoretical molecular construct is shown here (Fig 2e). We observe a bimodal distribution in both assays, and the height of the two peaks from the 5’ assay was comparable (Fig 2f), whereas, for the 3’ assay, one peak was visibly lower than the other (Fig 2g) and for both assays, the two major peaks were contributed by different strands. Here, the forward strand refers to reads with polyA/TSO and barcode detected in the original sequence, and the reverse strand refers to when the elements are found in the reverse complement. The mode of the barcode start-position is 90 for the forward strand and 78 for the reverse strand in the 3’ assay (Fig 2f) and the 5’ assay (Fig 2g). For the 3’assay reads, the mode of barcode start-position is 118 and 106 for the forward and reverse strands, respectively, with lower counts in the forward strand (Fig 2f). For the 5’assay reads, the mode of TSO start-position is 116 and 104 for forward and reverse strands, respectively (Fig 2g). The 2bp difference between the 5’ and 3’ assays (116 vs. 118 and 104 vs. 106), is likely caused by the difference in UMI length (10 vs. 12bp in the two assays). Taken together, the start position detected in the forward strand is 12 bases downstream for both assays compared to those detected in the reverse strand. However, a bias against polyA/barcoded detecting in the forward strand is only present in the 3’ assay reads. Importantly, in other in-house replicates, we found the difference between the two peak heights to be present in both the 5’ and 3’ assay for one batch of sequencing and not present in the other batch of the same sample (Fig S2e). Thus, whether a strand is sequenced preferentially appears to be variable for both assays. We then calculated the average Phred score of the first 200 bps of each read where the polyA for 3’ assay and TSO for 5’ assay is detected. We found that the distribution of the average Phred score for barcoded reads is similar between both kits (Fig 2h), which corresponds to the similar barcode identification rate between assays (Fig 2d). When separating the reads by strand, we found the forward strands in the 3’ assay have lower average Phred scores compared to the reverse strand, while both strands had similar average Phred scores in the 5’ assay (Fig 2i). This may contribute to the bias against polyA/barcoded detecting in the forward strand in the 3’ assay in this experiment.

### Fidelity of long-read description between 3’ and 5’ assay

Understanding to which extent both assays can generate complete cDNA molecules is fundamental for an accurate analysis of single-cell isoforms. To investigate this, we first looked at the long-read length distribution for each gene separately. Overall, average read length was comparable for most genes, but some of them were found to be longer in either the 3’ assay (n=44) or the 5’ assay (n=822) (Fig 3a). Since most of these genes associated with longer reads show differences in the 5’ assay, we investigated them further to understand whether this outlier group possesses distinct properties causing its outlier behavior. We considered the genes that had cDNAs over 500 bases longer in the 5’ assay compared to the 3’ assay and termed them the “5’assay-longer” (13.69%, n=822) while referring to the remaining genes with broadly similar cDNA lengths as the “5’-3’-similar genes”. We find the 5’assay-longer genes to have a longer average transcript length per gene in the annotation (p < 10^−10^, Wilcoxon Rank-Sum test) (Fig 3b). We found no differences between both gene groups in average intron length per gene (Fig S3a), but the 5’-3’-similar genes tend to have more introns per gene than the both-assay-similar genes (p < 10^−10^, Wilcoxon Rank-Sum test) (Fig S3b). In order to define whether the cDNAs captured by each assay covered full transcript annotation, we then investigated if the completeness of captured reads differed. The “read completeness” is calculated as the percentage of reads with extremities matching to both a transcription start site (TSS) and a polyA site for each gene in both libraries. Although the difference is for “5’-longer” genes is much stronger than for “5’-3’-similar” genes, the 5’ assay showed a significantly higher read completeness rate for both groups of genes compare to the 3’ assay (Fig. 3c). Interestingly, most of this can be attributed to the reads reaching a TSS with the 5’ assay, for both the 5’-longer and 5’-3’-similar genes (Fig 3d), while the percentage of reads matching polyA sites showed no difference between the 3’ and 5’ assay for genes in 5’ assay-longer genes (Fig. 3e). Additionally, the 3’ assay showed a slightly but significant higher rate of polyA site matching for 5’-3’-similar genes (Fig. 3e). In order to visualize this on a case-by-case basis, we plotted reads from both assays with ScisorWiz (Stein et al., 2022). Amongst the 5’-longer genes, we chose the PABPC1 gene as an example, with over 100 reads in both libraries. Clearly, the reads from the 3’ assay have more truncated reads toward the 5’ end (right side) compared to the 3’ end (left side) of the gene annotation (Fig. 3g). Other examples of 5’-longer genes also showed similar truncation in reads from the 3’assay (Fig. S3c-e). Finally, we investigated whether there were differences in expression levels between the two groups. We find that gene expression for 5’-longer genes is significantly lower than for 5’-3’-similar genes in both assays (Fig. 3f). While the genes in the 5’-longer group showed a slightly higher expression in the 5’ compared to 3’ assay group in bulk, there is no difference in individual cell types. These findings were broadly replicated in all tested cell types (Fig. S4). In summary, for most genes, no major differences in read length and completeness can be found; however, for 13.7 % of genes, the 5’ assay allows for the detection of longer and more complete transcripts, especially amongst genes with lower expression levels. These findings are reproduced in a separate dataset where 3’ and 5’ assay is performed on an independent experiment with same PBMC sample used for both assays (Fig. S2).

**Fig. 2.**
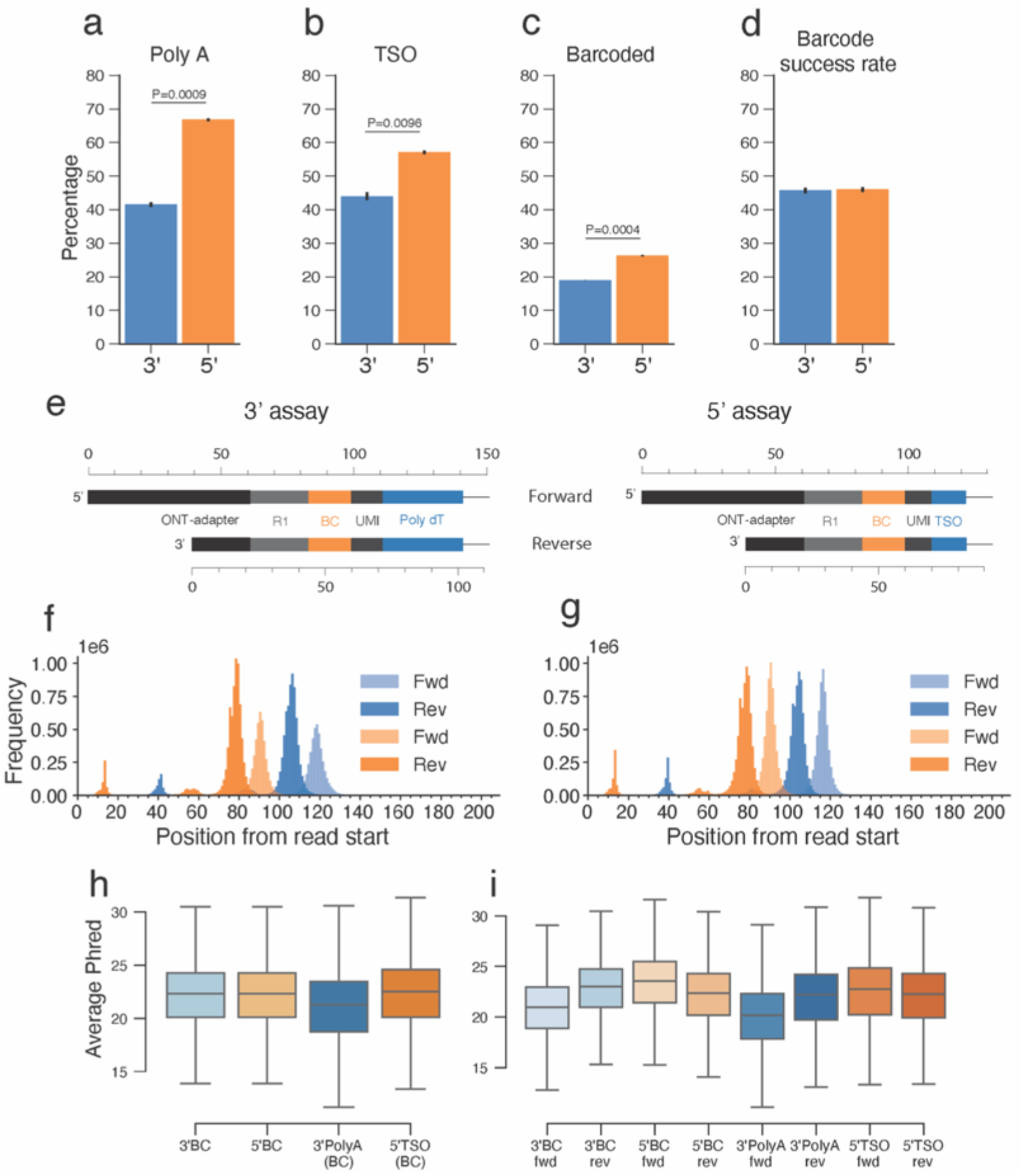
Barcode identification of single-cell 3’ and 5’ assay long-read sequencing. **a**. Barplot showing the percentage of read found with PolyA. **b**. Percentage of read found with TSO. **c**. Percentage of read found with barcodes given PolyA/TSO found for 3’ and 5’ assay respectively. **d**. Percentage of read found with barcode. *P*-values were calculated with the Student’s t-test. **e**. Schematic of the beginning of the theoretical molecular constructs. **f**. Histogram showing the distribution of start position of poly dT (blues) and 3’ assay barcodes (oranges) separated by strand. **g**. Histogram showing the distribution of start position of TSO (blues) and 5’ assay barcodes (oranges) separated by strand. Forward strand (Fwd) refers to reads with polyA/TSO and barcode detected in the original sequence, and the reverse strand (Rev) refers to when the elements are found in the reverse complement. **h**. Boxplot showing average Phred score of first 200 bp. BC: Barcoded reads. 3’PolyA: reads with poly dT from 3’ assay. 5’TSO: reads with TSO from 5’ assay. **i**. Boxplot showing average Phred score of first 200 bp separated by strand.

**Fig. 3.**
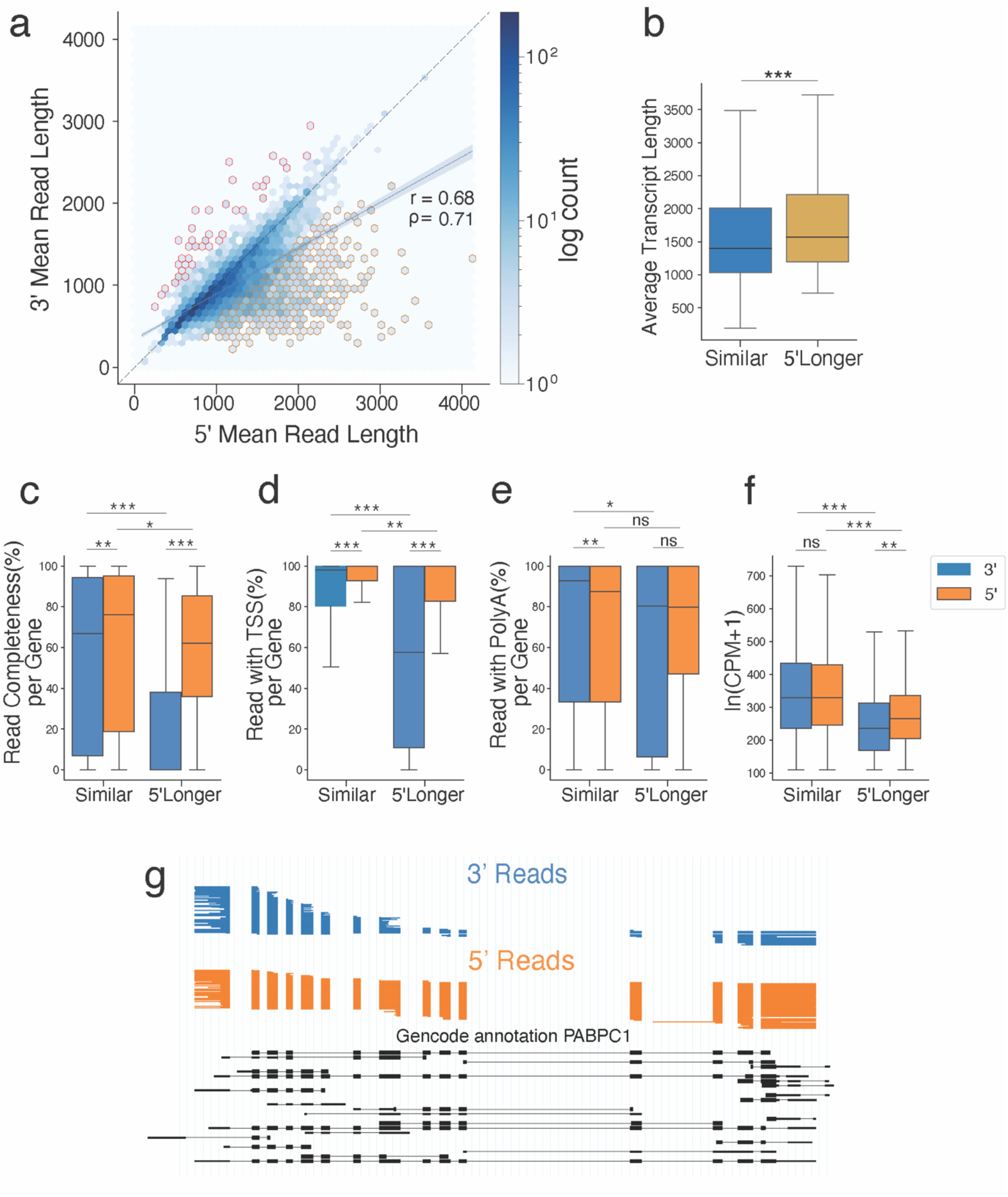
Fidelity of long-read description between 3’ and 5’ assay. **a**. Scatter plot of average read length per gene between 3’ and 5’ assay. Color scale showing the log count of genes. Regression line (blue) with 95% confidence interval in shaded blue bands. The yellow border represents 5’assay-longer genes (genes whose cDNAs are 500 bases longer in the 5’ assay compared to the 3’ assay; 13.69%, n=822). The red border represents genes longer in the 3’ assay with the same criteria (0.73%, n=44). Pearson’s r=0.69. Spearman’s rho=0.71. **b**. Boxplot showing the average transcript length per gene from the annotation of both-assay-similar genes (Similar) and 5’assay-longer genes (5’Longer) **c**. Boxplot of read completeness per gene of 3’ and 5’ assay comparing 5’assay-longer genes (5’Longer) and both-assay-similar genes (Similar). **d**,**e**. Boxplot of the percentage of read with TSS/polyA site per gene. **f**. Boxplot of the log-transformed counts-per-million (CPM) per gene. *P*-values were calculated using a two-sided Wilcoxon rank-sum test (***p < 10^−10^, **p < 10^−5^, *p < 0.05). **g**. cDNA reads captured by 3’ and 5’ assays for the PABPC1 gene. Each horizontal line indicates one read, colored by assays; clustered blocks denote exons. Black denotes annotated GENCODE transcripts.

### Comparison of cell-type-specific isoform expression

The advantage of single-cell long-read sequencing is to determine isoforms whose expression is distinct between different cell types. Thus, we have implemented differential isoform expression (DIE), TSS, Poly A site, and exon tests for single-cell (Gupta et al., 2018; Joglekar et al., 2021) and single-nuclei isoform sequencing (Hardwick et al., 2022). To assess if there are biases between the 3’ and 5’ assay in terms of the differential isoform testing between pairs of clusters, we picked the four largest clusters, namely Monocytes, CD4+ T cells, CD8+ T cells, and B cells, and considered all pairs of these clusters. We merged the two replicates from each assay and performed DIE tests on all pairs, once for the 5’ assay and once for the 3’ assay. We found an average of 116 significant genes for the six pairwise comparisons and only further considered those with over 100 significant genes. The number of genes tested significant between Monocytes and CD4+ T cells is the largest among all six comparisons, with 291 genes and 128 in both assays. 59 percent of the gene in the 3’ assay is confirmed by the 5’ assay, and 63 percent of the genes in the 5’ assay are confirmed by the 3’ assay (Fig. 4a). This leaves the question of whether the gene tested significant in one but not in the other assay are caused because of cut-off questions or the behaviors of the genes are fundamentally different between the assays. For each gene that tested significant in either assay, we choose the isoform with the largest delta PI between the two clusters and compare the delta PI value of the isoform from each assay. Overall, the dPI values from both assays correlate highly (Spearman correlation rho = 0.72, Pearson’s r = 0.77). Additionally, we observed few strongly contradictory genes because most genes (92%) fall into the lower left and upper right quadrants (Fig. 4b). We observed similar behaviors in the other cluster comparisons (Fig. 4c-f). We used a similar approach to test TSS and PolyA sites and find broadly similar results (Fig. S5). 57 percent of genes significant in the 3’ assay are confirmed by the 5’ assay, and 83 percent of significant genes in the 5’ assay is confirmed by the 3’ assay in the TSS test (Fig. S5a). Therefore, overall, we observed that 3’ and 5’ assays, despite the difference in read length in a small group of genes, have a similar profile of cell-type-specific isoforms.

**Fig. 4.**
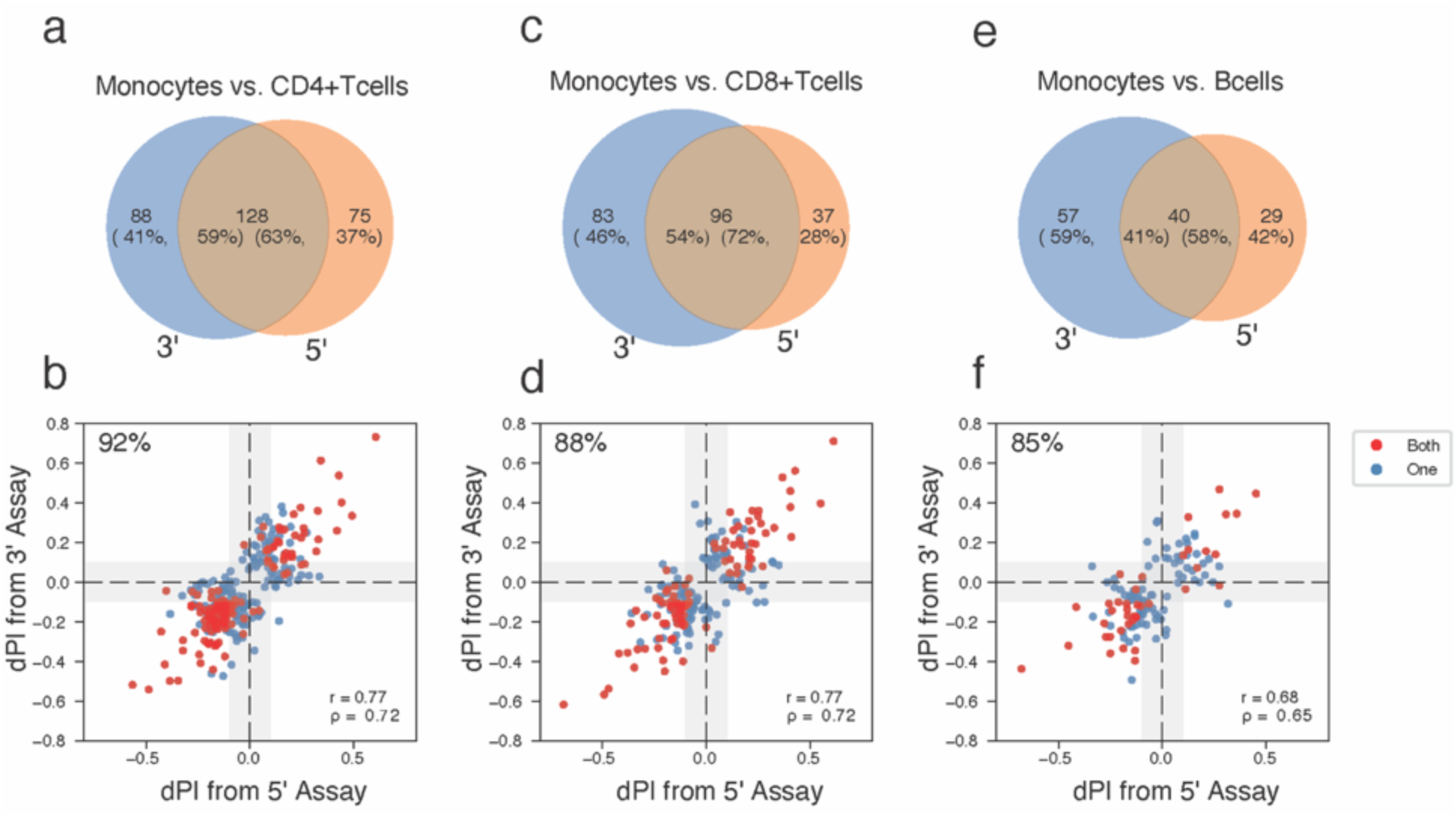
Comparison of cell-type-specific isoform expression. **a**. Venn diagram showing the number of genes tested significant with DIE between Monocytes and CD4+Tcells from 3’ and 5’ assay. **b**. Scatter plot showing the dPI value between 3’ and 5’ assay for genes testable in both and significant in either between Monocytes and CD4+Tcells. Isoform for each gene was chosen with the largest PI among the cell types and assays. Points are colored by whether the gene is tested significant in both assays or in one. Pearson’s r=0.72. Spearman’s rho=0.72. 92% of genes fall into the upper right and bottom left quadrants. **c**. Venn diagram showing the number of genes tested significant between Monocyte and CD8+Tcells in the same testing condition as a. **d**. Scatter plot showing the dPI value between 3’ and 5’ assay between Monocytes and CD8+Tcells in the same format as b. **e**. Venn diagram showing the number of genes tested significant between Monocyte and Bcells in the same testing condition as a. **f**. Scatter plot showing the dPI value between 3’ and 5’ assay between Monocytes and Bcells in the same format as b.

## Discussion

Barcoding strategies are fundamental to droplet-based single-cell sequencing, and understanding the biases and caveats between approaches is essential. Here, we comprehensively evaluated both short-read and long-read captured by the two marketed approaches from 10x Genomics, the “3’ assay” and the “5’ assay”. The two assays capture different ends of the transcript in the final library, while both use polydT primer for reverse transcription. Regardless of having different theoretical cDNA constructs, the barcode identification rate in the long-read is similar in both assays. Both assays have similar results in cell-type identification, and the gene expression profile is highly correlated in individual cell types. However, the 5’ assay captured more exonic UMIs and fewer intronic UMIs compared to the 3’ assay. It is possible that this could be due to the added barcode and UMI sequences that are located adjacently to poly(dT) primer in the 3’ assay – which is, molecularly speaking, the most obvious difference between the two assays.

The average barcoded read length of the 5’ assay is 127 bp longer than the 3’ assay. We found a group of genes captured with a longer average cDNA length per gene in the 5’ assay compared to the 3’ assay. This group of genes is lowly expressed and has a longer average transcript length than other expressed genes. The 5’ assay captures this group of genes with higher read completeness (the percentage of reads with both a known TSS and polyA identified) compared to the 3’ assay. The ability to preserve complete reads is essential in defining complete isoforms. Previous technical notes from 10x Genomics reported that the 3’ assay has a higher internal priming rate than the 5’ assay (“Technical Note - Interpreting Intronic and Antisense Reads in 10x Genomics Single Cell Gene Expression Data,” 2021). More incomplete reads and intronic reads could arise from the priming of poly dT at random A-rich regions in the 3’ assay. Therefore, for the investigation of the exon-combinatorics (Gupta et al., 2018; Hardwick et al., 2022; Tilgner et al., 2015; Tilgner et al., 2018), as well as single-nucleotide variants on exons, the 5’ assay offers advantages.

Despite the difference in read length in a small group of genes, cell-type-specific isoforms profiles observed from the two assays remain highly correlated. The 5’ assay has an advantage in investigating complete isoforms for genes that are lowly expressed and with longer average transcript lengths. Thus, similarly to the above observations, both assays perform similarly in terms of TSS and poly(A)-site assignments for most genes. However, for a subset of genes, which is characterized by high intron numbers and low expression overall, the 5’assay has strong advantages in TSS completeness with a slight disadvantage in poly(A) site - completeness.

One limitation of our current study is that the two assays are evaluated on different biological samples. We addressed this by additionally comparing both assays with another separate in-house sample and confirmed our main observations (Fig. S2). In summary, the 5’ and 3’ assays yield correlated results, but when TSS-choice or genes with very high intron numbers are of specific interest, the 5’ assay can offer advantages.

## Methods

### 10x Genomics 3’ and 5’ single-cell capture and Illumina library preparation

Human PBMCs (obtained from AllCells) were loaded on to the Chromium X instrument following the library preparation protocols in the Chromium Next GEM Single Cell 3’ HT Reagent Kits v3.1 User Guide (CG000416) and Chromium Next GEM Single Cell 5’ HT Reagent Kits v2 User Guide (CG000423), respectively. 3’ HT Libraries were sequenced on an Illumina NovaSeq with paired-end dual-indexing (28 cycles Read 1, 10 cycles i7, 10 cycles i5, 90 cycles Read 2). 5’ HT Libraries were sequenced on an Illumina NovaSeq with paired-end dual-indexing (26 cycles Read 1, 10 cycles i7, 10 cycles i5, 90 cycles Read 2).

### Alignment of single-cell short-read data

All of the 3′ and 5′ flowcells were demultiplexed with bcl2fastq (Illumina). FASTQ files were processed with Cell Ranger (10x Genomics), using the cellranger count pipeline on each GEM well with the GRCh38-2020-A reference to produce gene-barcode matrices and other output files, followed by aggregation of GEM wells with the cellranger aggr pipeline.

### Short-read data analysis

The 10x Cell Ranger output files were imported into Seurat (version 4.0.2) (Hao et al., 2021). For all samples, cells with less than 500 unique gene counts were removed for further analysis. The percentage of mitochondrial gene expression was regressed, and the gene expression matrix was normalized and scaled with Seurat’s “SCTransform” function. We used Seurat’s “merge” function to combine the replicates of each sample. We then clustered all cells with the Louvain algorithm using 15 principal components (PCs) and default resolution. We then annotate cells with SingleR and celldex (Aran et al., 2019) against the immune cell populations reference expression dataset (Consortium, 2012; Martens & Stunnenberg, 2013).

### PromethION library preparation and sequencing of long-read cDNA

For the first two technical replicates of each assay, 200 fmol cDNA was used to generate Oxford Nanopore Technology (ONT) libraries following the Amplicon barcoding with Native Barcoding Expansion 96 (EXP-NBD196, SQK-LSK109) protocol. The ONT libraries were then sequenced with PromethION flowcell (FLO-PRO002) for 72 hours. The two replicates were pooled and sequenced on one flowcell for each assay. Sequences were base called using Guppy (version 4.0.11) to generate fastq file.

### Barcode identification of 3’ and 5’ long read

For each read and its reverse complement, we located constant sequences next to the cellular barcode in the first 200 bases. For the 3’ assays, those are stretches of 9 consecutive Ts (T9), and for the 5’ assays, are the Template Switch Oligo (TSO) sequence. We then searched for perfectly matching 16-mer barcodes between the read start and polyA (3’ assay) or TSO (5’ assay). UMI sequences between the end of the barcode and the start of polyA (3’ assay) or TSO (5’ assay) are recorded for each read.

### Alignment and characterization of long reads

Filtered fastqs with barcoded long reads were mapped and aligned to the reference genome (hg38) using Minimap2 with the built-in function “MMalign” in scisorseqr. Scisorseqr’s “MapAndFilter” and “InfoPerLongRead” functions were used to filter the aligned reads and denote the completeness of each read using published CAGE and polyA site data (Herrmann et al., 2020; Lizio et al., 2015). The full procedure was described in our previous publication and packaged into scisorseqr’s workflow (Joglekar et al., 2021). Technical replicates were pooled for long-read analysis. To factor out PCR duplication, one transcript per molecule (barcode+UMI+gene) was chosen for analysis. The sample with a higher number of transcripts was randomly down-sampled to the same size as the other.

### Differential isoform, polyA, and TSS tests

The quantifying of isoform and testing details were described in our previous publication (Joglekar et al., 2021), and packaged into scisorseqr’s “IsoQuant” and “DiffSplicingAnalysis” functions, but we outline the process here as well. Each read is represented by a string denoting the TSS, introns, and polyA-site, symbolizing an isoform. An ID number is assigned to each present isoform, from the most abundant to the least. Counts of each isoform ID were assigned to each cell type. For each differential test between two cell types, a maximum of an 11×2 matrix of isoform counts x cell-type was constructed for genes with sufficient depth (25 reads/gene in cell-type). The first ten rows of the 11×2 matrix correspond to the top ten abundant isoforms, and the last row is the counts of all the other isoforms if present. For each gene, *p*-values from the *X*^*2*^ test were reported, and the ΔPI value (the sum of absolute change in percent isoform (PI) of the top two isoforms) was calculated. Benjamini Hochberg (Benjamini & Hochberg, 1995) correction for multiple testing was applied to return a corrected p-value with a false discovery rate of 5%. The gene was considered to be significantly differentially spiced if the FDR corrected p-value was <= 0.05 and ΔPI greater than 0.1. For differential ployA or TSS tests, polyA sites or TSS are determined for each read. The counts of TSS or polyA were summarized and reduced into an 11×2 table as described above. Testing was performed as described above for isoforms.

