## Supplementary figures for "Comparing 10x Genomics single-cell 3’ and 5’ assay in short-and long-read sequencing"

### Contents

Supplementary Figure 1: Short read analysis of single-cell 3' and 5' assays with PBMC data

Supplementary Figure 2: Analysis replicated with a single PBMC sample for both assays

Supplementary Figure 3: Fidelity of long-read description between 3' and 5' assay

Supplementary Figure 4: Fidelity of long-read description between 3' and 5' assay per cell type

Supplementary Figure 5: Comparison of cell-type-specific TSS and polyA-site expression

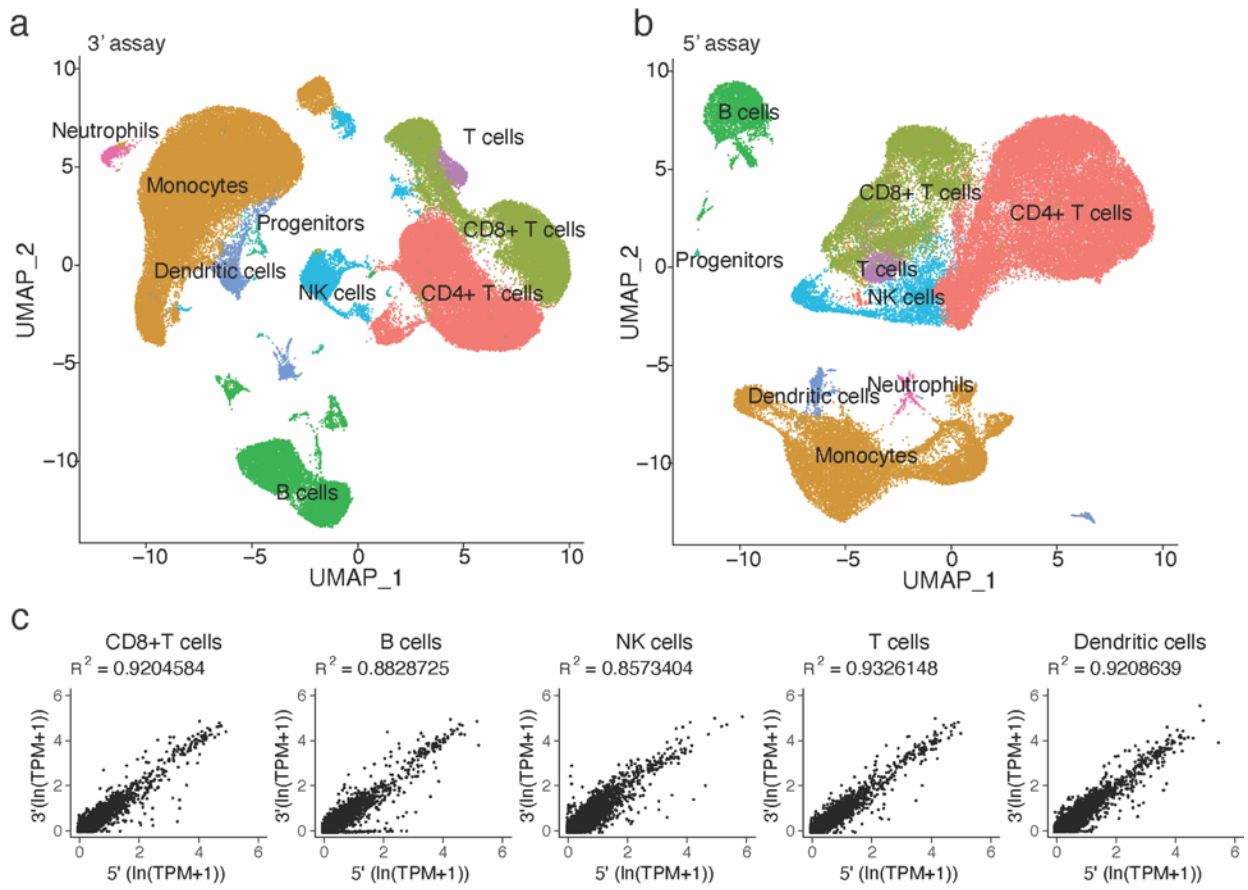


**Supplementary Figure 1: Short read analysis of single-cell 3' and 5' assays with PBMC data**

**a.** UMAP of PBMC data from 3' assay. **b.** UMAP of PBMC data from 5' assay. Cell types identified by SingleR package. **c.** Scatter plot showing the gene expression level of the same cell types between 3' and 5' assay.

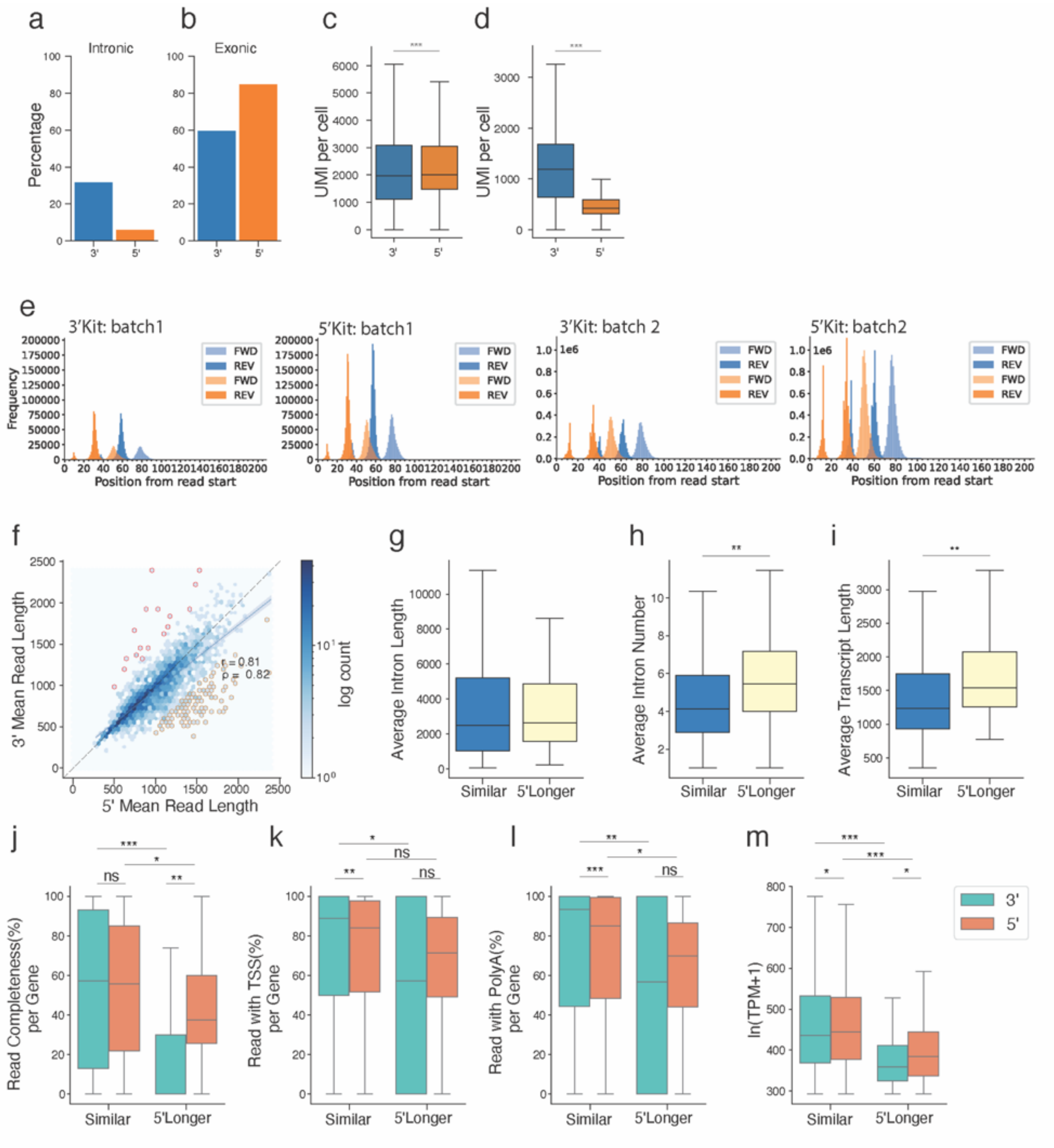


**Supplementary Figure 2: Analysis replicated with a single PBMC sample for both assays** **a.** Barplots showing the average percentage of mapped reads in introns and **b.** the average percentage of mapped reads in exons of two replicates from both assays. **c.** Boxplots of exonic UMI per cell, **and d.** boxplots of intronic UMI per cell. Down-sampled each replicate to 10 thousand reads per cell after excluding reads with no barcode and reads for cells with less than 10 thousand reads. **e.** Histogram showing the distribution of start position from two batches of sequencing. Color-coded by poly dT (blues) and 3' assay barcodes (oranges) for 3' assay and TSO (blues) and 5' assay barcodes (oranges) for 5' assay. Forward strand (Fwd) refers to reads with polyA/ TSO and barcode detected in the original sequence, and the reverse strand (Rev)

refers to when the elements are found in the reverse complement. **f.** Scatter plot of average read length per gene between 3' and 5' assay. Color scale showing the log count of genes. Regression line (blue) with 95% confidence interval in shaded blue bands. The yellow border represents 5'assay-longer genes (genes whose cDNAs are 500 bases longer in the 5' assay compared to the 3' assay). The red border represents genes longer in the 3' assay with the same criteria. Pearson's r=0.81. Spearman's rho=0.82. **g.** Boxplot showing the average intron length per gene from the annotation of both-assay-similar genes (Similar) and 5'assay-longer genes (5'Longer). **h.** Boxplot showing the average intron numbers per gene from the annotation of both-assay-similar genes (Similar) and 5'assay-longer genes (5'Longer). **i.** Boxplot showing the average transcript length per gene from the annotation of both-assay-similar genes (Similar) and 5'assay-longer genes (5'Longer). **j.** Boxplot of read completeness per gene of 3' and 5' assay comparing 5'kit-longer genes (5'Longer) and both-kit-similar genes (Similar). **k,l.** Boxplot of the percentage of read with TSS/ polyA site per gene. **m.** Boxplot of the log-transformed counts-per-million (CPM) per gene. *P*-values were calculated using a two-sided Wilcoxon rank-sum test (***$p < 10^{-10}$, **$p < 10^{-5}$, *$p < 0.05$).

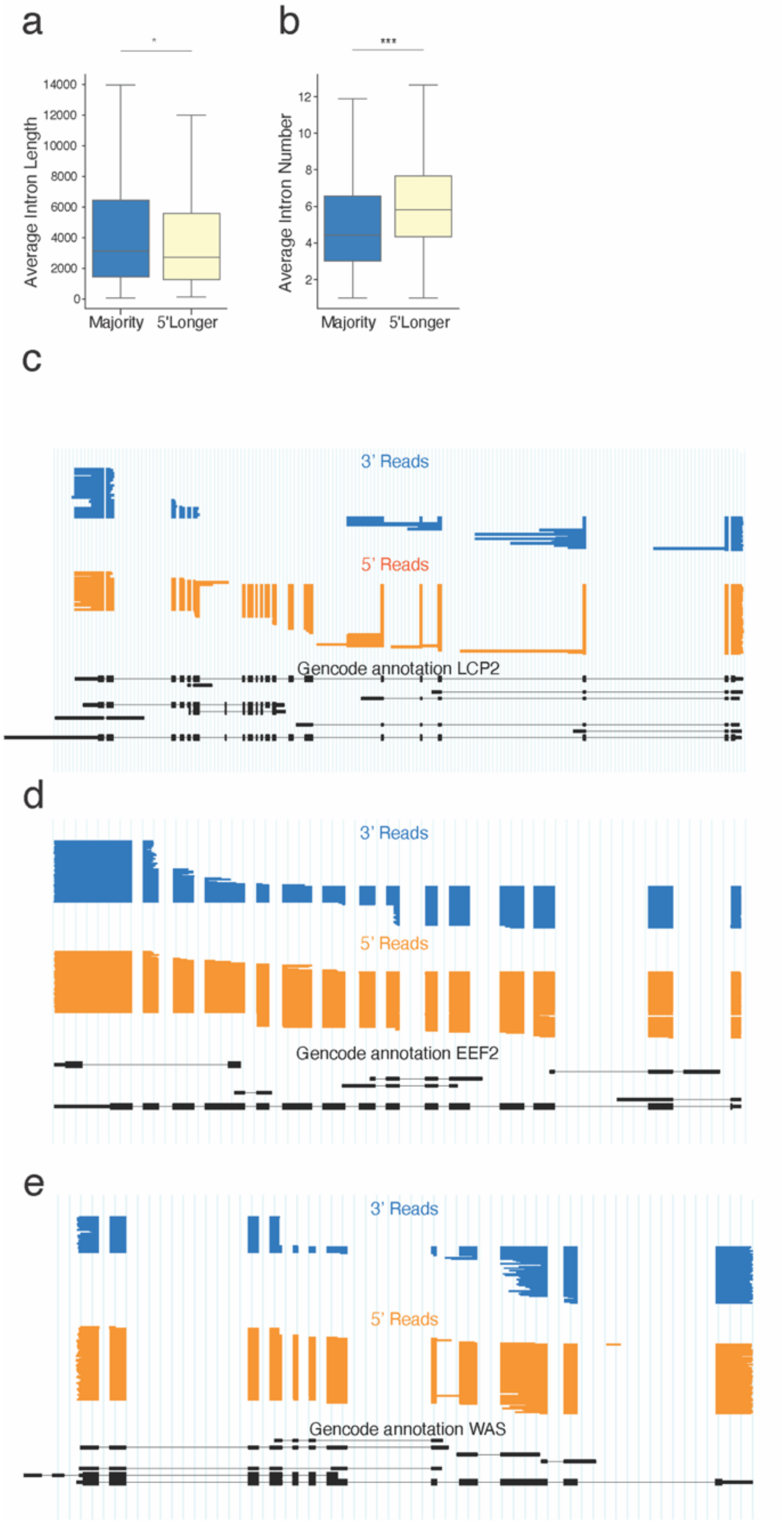
a
b
Average Intron Length
Average Intron Number
Majority
5'Longer
c
3' Reads
5' Reads
Gencode annotation LCP2
d
3' Reads
5' Reads
Gencode annotation EEF2
e
3' Reads
5' Reads
Gencode annotation WAS

**Supplementary Figure 3: Fidelity of long-read description between 3' and 5' assay**
**a.** Boxplot showing the average intron length per gene from the annotation of both-assay-similar genes (Similar) and 5'assay-longer genes (5'Longer). **b.** Boxplot showing the average intron numbers per gene from the annotation of both-assay-similar genes (Similar) and 5'assay-longer genes (5'Longer). **c.** cDNA reads captured by 3' and 5' assays for the LCP2 gene. Each horizontal line indicates one read, colored by assays; clustered blocks denote exons. Black denotes annotated GENCODE transcripts. **d.** cDNA reads captured by 3' and 5' assays for the EEF2 gene. **e.** cDNA reads captured by 3' and 5' assays for the WAS gene.

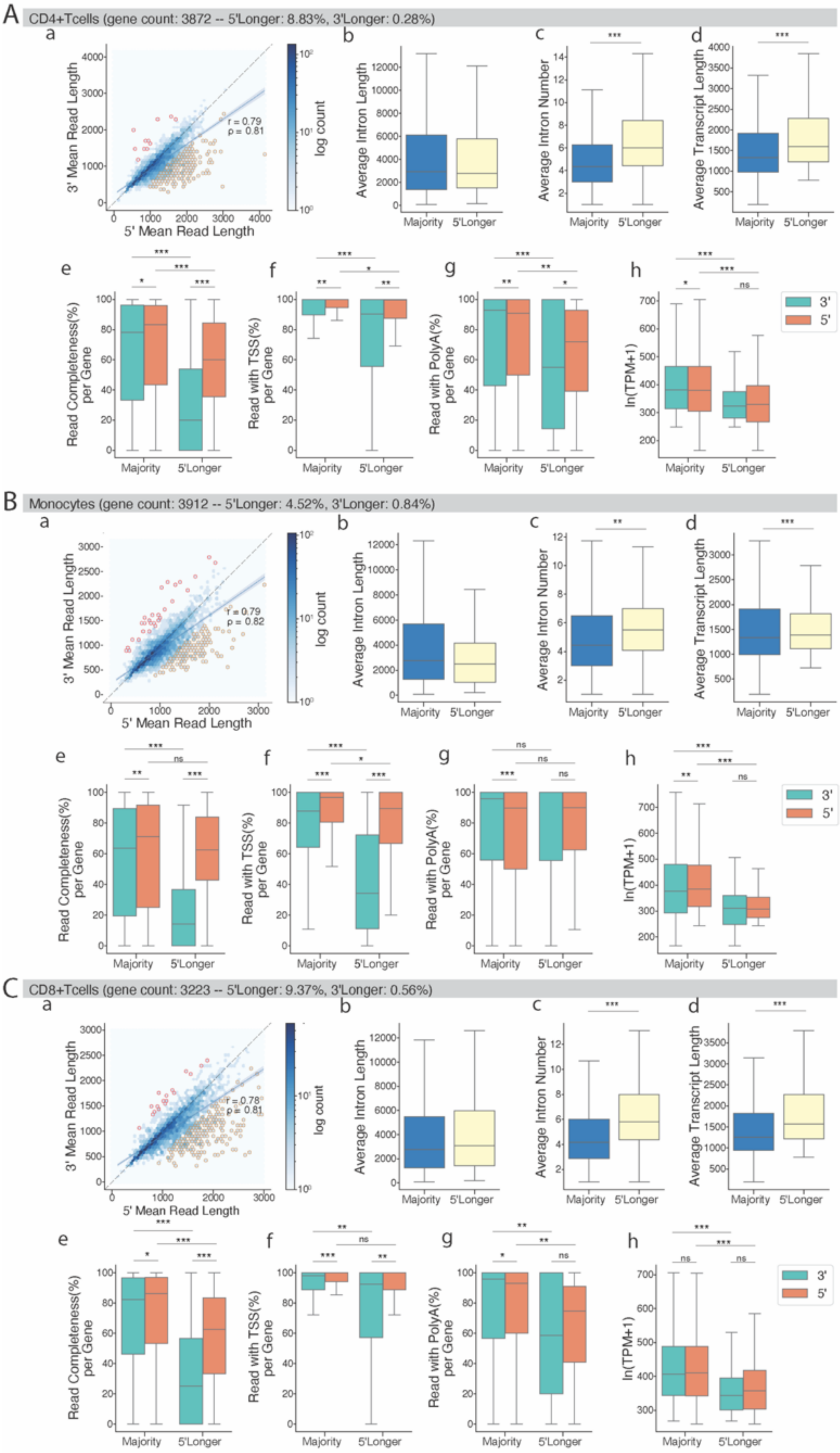
A
CD4+Tcells (gene count: 3872 -- 5'Longer: 8.83%, 3'Longer: 0.28%)
3' Mean Read Length
5' Mean Read Length
r = 0.79
ρ = 0.81
log count
Average Intron Length
Average Intron Number
Average Transcript Length
Majority
5'Longer
Read Completeness(%) per Gene
Read with TSS(%) per Gene
Read with PolyA(%) per Gene
ln(TPM+1)
3'
5'
B
Monocytes (gene count: 3912 -- 5'Longer: 4.52%, 3'Longer: 0.84%)
r = 0.79
ρ = 0.82
C
CD8+Tcells (gene count: 3223 -- 5'Longer: 9.37%, 3'Longer: 0.56%)
r = 0.78
ρ = 0.81

**Supplementary Figure 4: Fidelity of long-read description between 3' and 5' assay per cell type**

Large panels are the results for each cell type**: A.** CD4+Tcells, **B.** Monocytes, **and C**. CD8+Tcells. Small panels are the same for each cell type which is as follows, **a.** Scatter plot of average read length per gene between 3' and 5' assay. Color scale showing the log count of genes. Regression line (blue) with 95% confidence interval in shaded blue bands. The yellow border represents 5'assay-longer genes (genes whose cDNAs are 500 bases longer in the 5' assay compared to the 3' assay). The red border represents genes longer in the 3' assay with the same criteria. **b.** Boxplot showing the average intron length per gene from the annotation of both-assay-similar genes (Similar) and 5'assay-longer genes (5'Longer). **c.** Boxplot showing the average intron numbers per gene from the annotation of both-assay-similar genes (Similar) and 5'assay-longer genes (5'Longer). **d.** Boxplot showing the average transcript length per gene from the annotation of both-assay-similar genes (Similar) and 5'assay-longer genes (5'Longer). **e.** Boxplot of read completeness per gene of 3' and 5' assay comparing 5'kit-longer genes (5'Longer) and both-kit-similar genes (Similar). **f, g.** Boxplot of the percentage of read with TSS/ polyA site per gene. **h.** Boxplot of the log-transformed counts-per-million (CPM) per gene. *P*-values were calculated using a two-sided Wilcoxon rank-sum test (***$p < 10^{-10}$, **$p < 10^{-5}$, *$p < 0.05$).

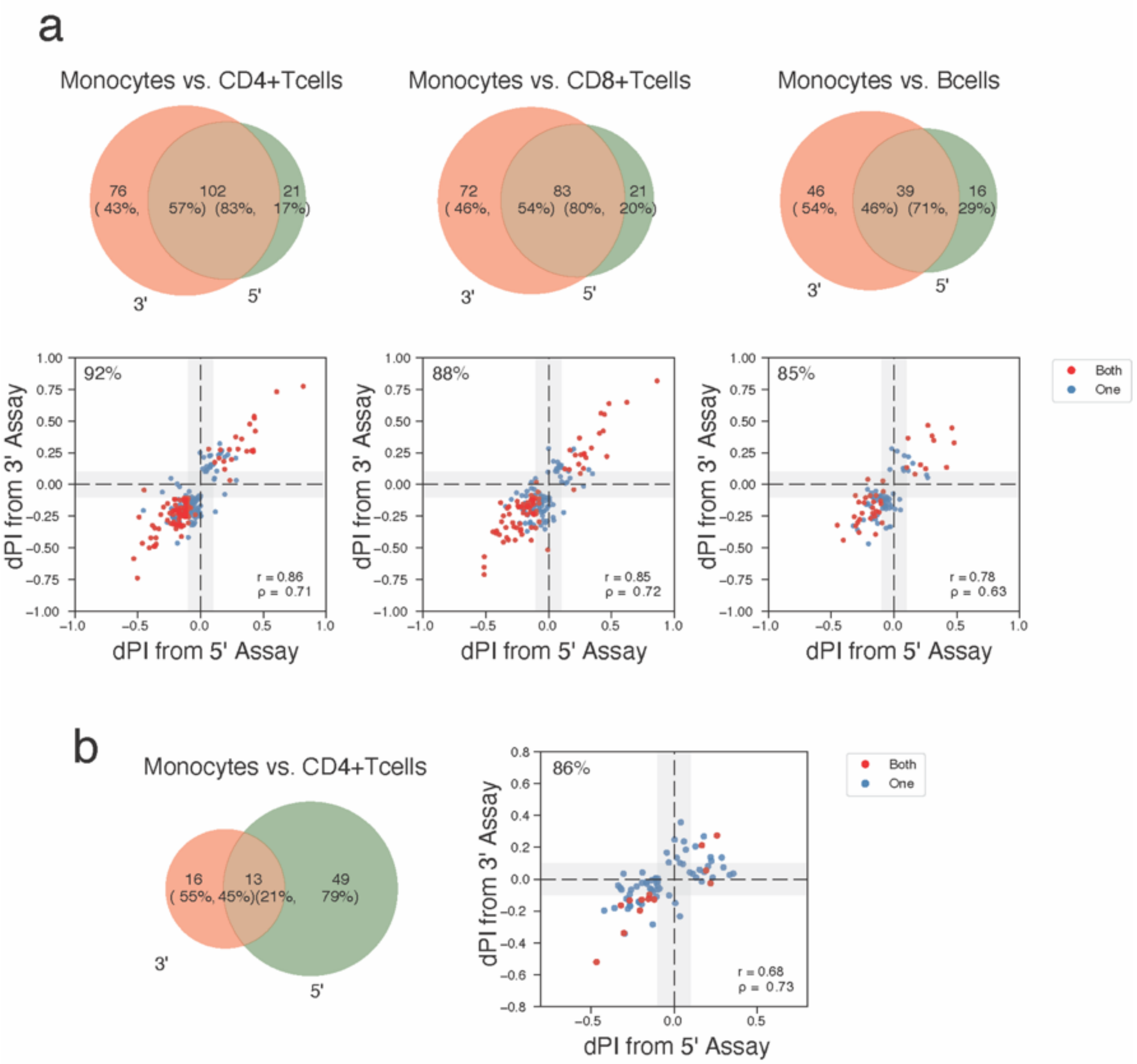


**Supplementary Figure 5: Comparison of cell-type-specific TSS and polyA-site expression**

**a.** Venn diagrams showing the number of genes tested significant with the differential TSS test between different cell types from 3' and 5' assay. Scatter plots showing the dPI value between 3' and 5' assay for genes testable in both and significant in either cell type. TSS for each gene was chosen with the largest PI among the cell types and assays. Points are colored by whether the gene is tested significant in both assays or in one. **b.** Venn diagrams showing the number of genes tested significant with the differential polyA-site test between Monocytes and CD4+Tcells from 3' and 5' assay. Scatter plots showing the dPI value between 3' and 5' assay for genes testable in both and significant in either Monocytes and CD4+Tcells. PolyA-site for each gene was chosen with the largest PI among the cell types and assays. Points are colored by whether the gene is tested significant in both assays or in one. Tests with less than fifty genes tested significant in either cell type in total are excluded.
